# GAPVD1 regulates CSNK1D-dependent PER2 FASP phosphorylation

**DOI:** 10.64898/2026.05.29.728667

**Authors:** Hussam Ibrahim, Yuhong Chen, Christina Buschhaus, Judith Düring, Lukas Lubczyk, Vivienne Ogasa, Sandra Szczutkowski, Ellen Wannagat, Hilal Watad, Viviane Zimmer, Christian Preisinger, Faiza Kalfalah, Hans Reinke

**Affiliations:** University of Düsseldorf, Medical Faculty, Institute of Clinical Chemistry and Laboratory Diagnostics, Düsseldorf, Germany; Interdisciplinary Centre for Clinical Research (IZKF), University Hospital RWTH Aachen, Aachen, Germany

## Abstract

The mammalian circadian clock relies on precisely timed transcriptional and post-translational events to generate ∼24-hour rhythms. A key post-translational mechanism is phosphorylation of PER2 by CSNK1D, governed by a phosphoswitch that integrates stabilizing and destabilizing phosphorylation marks at the FASP and Degron sites to control PER2 stability and circadian period. Here, a novel role for the PER complex protein GAPVD1 in regulating CSNK1D-mediated FASP site phosphorylation is revealed. GAPVD1 associates with PER2 and CSNK1D via a distinct helical bundle domain, weakening PER2-CSNK1D association and attenuating PER2 FASP phosphorylation in cells and in vitro. Mechanistically, this helical bundle interacts with the PER2 casein kinase binding domain and is contacted by the adjacent VPS9 region in GAPVD1, providing a structural explanation for how flanking GAPVD1 domains can control access of binding proteins to the helical bundle. Notably, a minimal GAPVD1 helical bundle fragment is sufficient for binding to PER2 and CSNK1D but not for robust attenuation of PER2 FASP phosphorylation, indicating that additional regions of GAPVD1 are required. GAPVD1-mediated inhibition of PER2 FASP phosphorylation also turned out to be independent of the CSNK1D C-terminal tail, pointing to mechanisms such as displacement of PER2 FASP from the CSNK1D active site or allosteric regulation of CSNK1D substrate engagement. Together, these findings identify GAPVD1 as a PER-complex scaffold that modulates the PER2-CSNK1D control node and introduce an additional regulatory layer into the circadian phosphoswitch.

A GAPVD1 helical bundle domain binds the PER2 CKBD and CSNK1D to weaken PER2-CSNK1D interaction and attenuate PER2 FASP phosphorylation.

## Introduction

The mammalian circadian clock is an intrinsic timekeeping system that drives robust biochemical, physiological, and behavioural rhythms with a period of approximately 24 hours. At its core, the clock operates via interconnected transcription-translation feedback loops in which a heterodimer of the transcriptional activators Circadian locomotor output cycles kaput (CLOCK) and Aryl hydrocarbon receptor nuclear translocator like (ARNTL/BMAL1) induces the expression of Period (PER) and Cryptochrome (CRY) genes, whose protein products assemble into a large macromolecular complex in the cytoplasm. Upon nuclear entry, the PER complex inhibits the transactivator function of CLOCK-BMAL1, thereby repressing its own transcription and closing the negative feedback loop (*1*). CRY proteins repress CLOCK-BMAL1 transcription by binding to the E-box-bound complex, while PER2 recruits Casein kinase 1 delta (CSNK1D), which phosphorylates the basic helix-loop-helix regions of CLOCK and BMAL1, reducing their DNA binding affinity. These events drive efficient removal of CLOCK-BMAL1 from gene promoters. Complex formation, cyclical phosphorylation, and targeted degradation of core clock proteins underlie the periodicity and stability of the circadian oscillator, ensuring precise temporal regulation of downstream physiological processes (*2*).

Phosphorylation of PER2 by CSNK1D is a central post-translational modification step that governs the dynamics of the clock through the so-called phosphoswitch mechanism. CSNK1D-mediated phosphorylation at two key regions of PER2, the FASP (familial advanced sleep phase) and Degron sites, determines PER2’s degradation rate via recruitment of the Beta-transducin repeat containing E3 ubiquitin protein ligase (BTRC), thereby controlling the key parameters of the circadian clock period length, entrainment, and temperature compensation (*3*–*6*). The phosphoswitch arises from hierarchical and competitive phosphorylation events, whereby stabilizing phosphorylation marks at the FASP site antagonize Degron phosphorylation and promote protein stability, while phosphorylation at the Degron site targets PER2 for proteasomal degradation. Regulation of CSNK1D activity through autophosphorylation, substrate competition, and conformational changes confers temporal precision and adaptability to the circadian clock, and is an active area of investigation (*7*).

Recent structural and biochemical studies have revealed that the regulatory C-terminal tail of CSNK1D can undergo dynamic autophosphorylation, acting as an inhibitory pseudosubstrate that modulates kinase activity and substrate specificity. This mechanism allows CSNK1D to tune its activity according to regulatory context and interaction with clock proteins. The precise timing and extent to which autophosphorylation of the C-terminal tail is required for regulating CSNK1D activity remain incompletely defined. Moreover, the selective interaction network of CSNK1D with its own tail and with partner or substrate proteins, which can likewise exert inhibitory effects through regulatory patches near the active site, is critical for controlling phosphorylation status, localization, and protein stability, with direct consequences for clock parameters and overall robustness of the oscillator (*8*).

GAPVD1 (GTPase-activating protein and VPS9 domains 1) has recently emerged as a component of the mammalian PER complex with dual roles in circadian timing and membrane trafficking (*9*–*11*). Proteomic and biochemical studies indicate that GAPVD1 interacts most closely with PER2 and CSNK1D. Notably, PER2 and CSNK1D regulate GAPVD1 phosphorylation, triggering GAPVD1 degradation (*11*). Whether GAPVD1 can reciprocally influence the phosphorylation dynamics of PER2, and by extension the operation of the phosphoswitch, has not been explored.

Given the centrality of the phosphoswitch in dictating core oscillator function, we set out to determine the impact of GAPVD1 on the PER2-CSNK1D interaction. By dissecting binding interfaces and phosphorylation states of PER2 complexes with GAPVD1 and CSNK1D, we identified a helical bundle region in GAPVD1 that interacts with the casein kinase binding domain in PER2 and with CSNK1D. Interaction with GAPVD1 weakens PER2-CSNK1D binding and leads to reduced phosphorylation of the PER2 FASP site. Mapping and mutational analyses further reveal that this same helical bundle is contacted by the adjacent VPS9 region in GAPVD1, providing a structural basis for how flanking GAPVD1 regions can influence PER2 binding and helical bundle phosphorylation, likely through intramolecular interactions. Notably, a minimal helical bundle fragment is sufficient for binding to PER2 and CSNK1D but fails to fully attenuate PER2 FASP phosphorylation, indicating that additional regions of GAPVD1 are required and pointing to mechanisms such as altered CSNK1D engagement with the FASP site. Our results reveal GAPVD1 as a novel regulator of CSNK1D-dependent PER2 phosphorylation and an additional layer of regulation within the molecular circadian clock.

## Results

GAPVD1 is a component of the cytoplasmic PER complex, and its knockdown leads to circadian period lengthening in mouse fibroblasts, indicating an essential role in maintaining normal oscillator function (*9*). Despite this, the molecular mechanisms behind GAPVD1-dependent regulation remain unclear. PER2 can interact with a variety of proteins, and both the quality and quantity of these interactions are themselves regulated and play key roles in shaping circadian clock function. PER2 protein stability, which is crucial for most functions of the cellular oscillator, is controlled by the phosphoswitch through phosphorylation by CSNK1D (*5*). Additionally, interaction of CRY proteins with PER2 blocks their FBXL3 (F-Box and Leucine rich repeat protein 3)-mediated degradation, while contacts of PER2 with nuclear receptors create additional molecular switches under clock control (*12*). Given this complexity, it is conceivable that GAPVD1 might modulate PER2 function by altering its posttranslational modifications or protein-protein interactions. Elucidating the physical contact points within the GAPVD1-PER2-CSNK1D complex might provide first insights into how GAPVD1 regulates PER complex function and circadian timing.

### A helical bundle in GAPVD1 binds PER2 and CSNK1D

First, we analysed a set of GAPVD1 deletion mutants for PER2- and CSNK1D-binding. The putative domain structure of GAPVD1 according to the Alphafold algorithm was inspected (*13*) (**Fig. 1A**), and GAPVD1 deletions were designed that supposedly do not disrupt stably folded subdomains (**Fig. 1B**). Full-length GAPVD1-tRFP (1-1478) and various GAPVD1 deletion mutants were co-expressed with PER2-GFP in HEK293T cells. Crude mapping showed that compared to the tRFP control the N-terminal half GAPVD1(1-690), which includes the RasGAP domain and a putative long alpha-helical stretch immediately N-terminal of it, does not significantly bind to PER2 or CSNK1D; neither does GAPVD1(Δ361-1367) comprising RasGAP and the other small GTPase regulating domain VPS9 (**Fig. 1C**). In contrast, interaction of GAPVD1(680-1478) and GAPVD1(353-1372) with PER2 and CSNK1D indicated that amino acids 680-1372 contain an interaction surface for binding to both proteins. Surprisingly, full-length GAPVD1 bound to CSNK1D but only very weakly to PER2, indicating that parts of the N-terminal half or the C-terminus of GAPVD1 might impede interaction specifically with PER2 (**Fig. 1C**).

**Figure 1.**
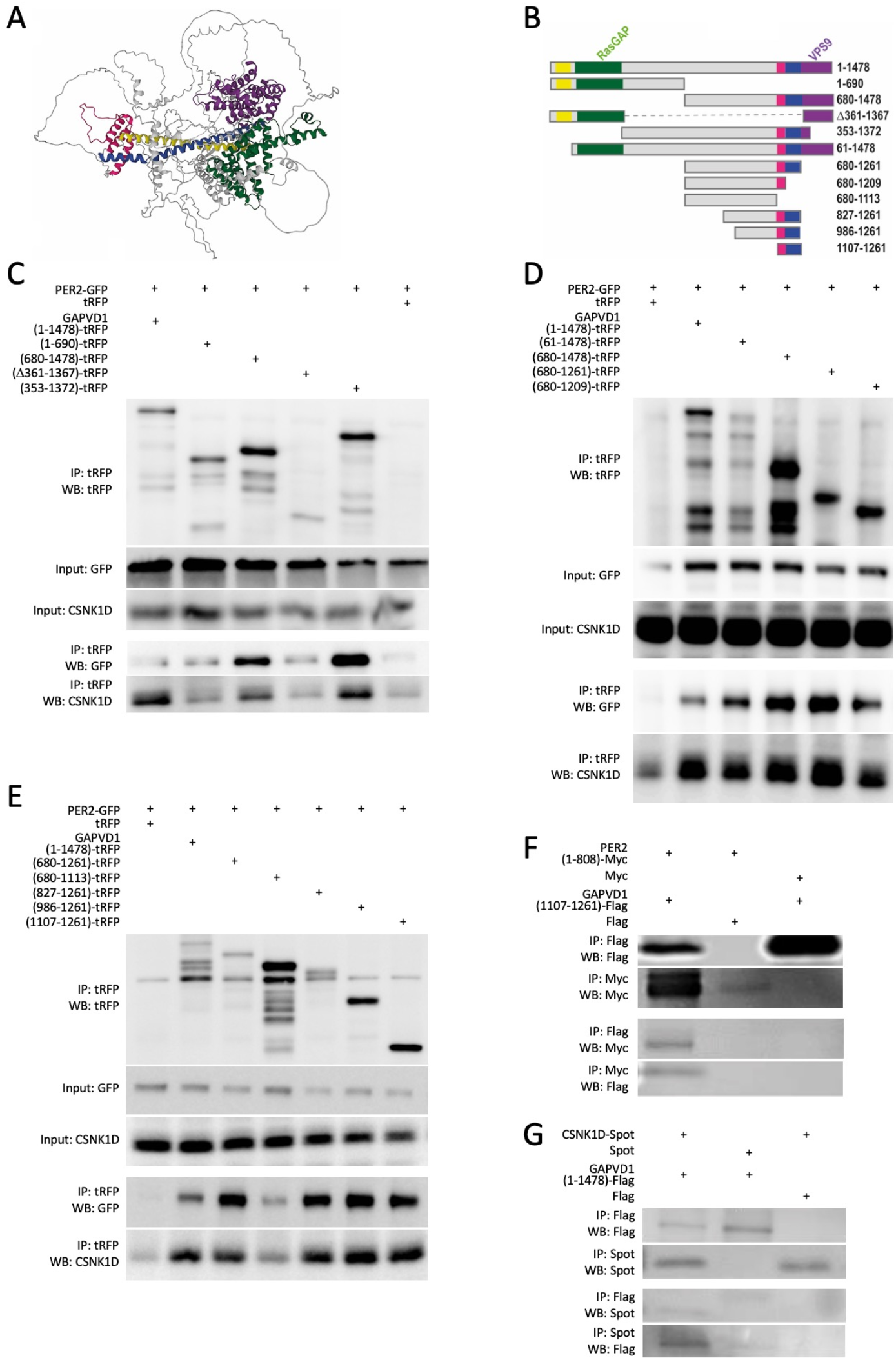
Mapping of a helical bundle domain in GAPVD1 that mediates binding to PER2 and CSNK1D. (**A**) Putative protein domains were determined by inspection of the human GAPVD1 structure proposed by Alphafold (*13*): Helical (1-60, yellow), RasGAP (61-446, green), IDR (447-1113, grey), Helical bundle (1114-1192, magenta), Helical (1193-1260, blue) and VPS9 (1335-1478, violet). (**B**) Schematic representation of human GAPVD1 domain organization and the C-terminally tRFP-tagged deletion constructs (including full-length GAPVD1(1–1478) and N- and C-terminal truncations) designed based on the Alphafold-predicted structure. (**C–E**) HEK293T cells were transiently transfected with PER2-GFP and tRFP or the indicated GAPVD1-tRFP constructs. Cell lysates were analyzed by immunoblotting and subjected to immunoprecipitation (IP) with anti-tRFP; IPs and inputs were probed with antibodies against tRFP, GFP, and CSNK1D. tRFP alone was used as control for non-specific IP. (**F**) In a Tobacco BY-2 cell-free expression system, GAPVD1(1107-1261)-Flag and PER2(1-808)-Myc were co-expressed, followed by anti-Flag or anti-Myc IP and immunoblot detection of Flag and Myc. (**G**) BY-2 lysates co-expressing GAPVD1(1-1478)-Flag and CSNK1D-Spot were subjected to reciprocal IP with anti-Flag or anti-Spot and analyzed by immunoblotting for Flag and Spot epitopes. In some experiments, input samples could not be shown due to low protein abundance or antibodies unsuitable for input detection; in these cases, a direct immunoprecipitation is displayed instead.

C-terminal truncations of the binding-competent variant GAPVD1(680-1478) revealed that deletion of amino acids 1210-1478, which contain the C-terminal VPS9 domain and a second putative long alpha-helical stretch, does not negatively affect binding (**Fig. 1D**). Likewise, binding of GAPVD1(1107-1261) showed that the C-terminal part of the intrinsically disordered region (IDR) from amino acids 680 to 1106 does not seem to be part of the interaction surface (**Fig. 1E**). However, binding of GAPVD1(680-1113) to both PER2 and CSNK1D was strongly diminished (**Fig. 1E**). Since the slightly larger variant GAPVD1(680-1209) was found to bind both proteins (**Fig. 1D**), amino acids 1114-1209 seem to be essential for these interactions. According to Alphafold this part of the protein folds into a compact domain, in which three alpha helices form a tight bundle that is located between the IDR and a putative coiled coil domain, which is formed by the two stretched-out alpha helices N-terminal of the RasGAP and VPS9 domains (**Fig. 1A**).

The smallest binding-competent fragment in these experiments was GAPVD1(1107-1261), which also interacted with PER2(1-808) when the proteins were expressed in a heterologous cell-free lysate derived from Tobacco BY-2 cells (*14*) (**Fig. 1F**). In the same in vitro system full-length GAPVD1(1-1478) interacted with CSNK1D, indicating that both interactions of GAPVD1 with PER2 and CSNK1D do not depend on other mammalian PER complex components (**Fig. 1G**).

### The helical bundle is a VPS9-regulated protein interaction module in GAPVD1

Mapping of the GAPVD1-PER2 interface showed that both removal of the N-terminal half of GAPVD1 and deletion of the C-terminal VPS9 domain increased binding of helical bundle-containing GAPVD1 variants to PER2, whereas binding to CSNK1D was not similarly enhanced **(Fig. 1C-E)**. This indicates that flanking regions may modulate access specifically to the PER2 interaction surface, e.g. through formation of intramolecular protein-protein contacts in GAPVD1. Because deletion of VPS9 involves one conserved regulatory module, it offered a structurally well-defined way to test this model, and we focused our subsequent analysis on this domain as a first step toward understanding how flanking regions might regulate access of binding proteins to the helical bundle. In BY-2 lysates, co-expression of the VPS9 domain as GAPVD1(1261-1478)-Myc with the minimal PER2 binding variant GAPVD1(1107-1261)-Flag yielded robust signals for both proteins after Flag immunoprecipitation, whereas replacing the Flag antibody with a GFP antibody in the immunoprecipitation abolished recovery of both proteins, confirming a specific interaction of the VPS9 domain with GAPVD1(1107-1261) **(Fig. 2A)**. Notably, GAPVD1(1261-1478) expressed alone as Myc- or Flag-tagged construct was barely detectable in the same assay, indicating that independently of the affinity tag the VPS9 domain is intrinsically unstable but is stabilized upon interaction with GAPVD1(1107-1261). Together, these observations indicate that the helical bundle and VPS9 regions can contact each other and support a model in which exposure of the GAPVD1 protein interaction domain is modulated by flanking regions through direct physical interactions.

**Figure 2.**
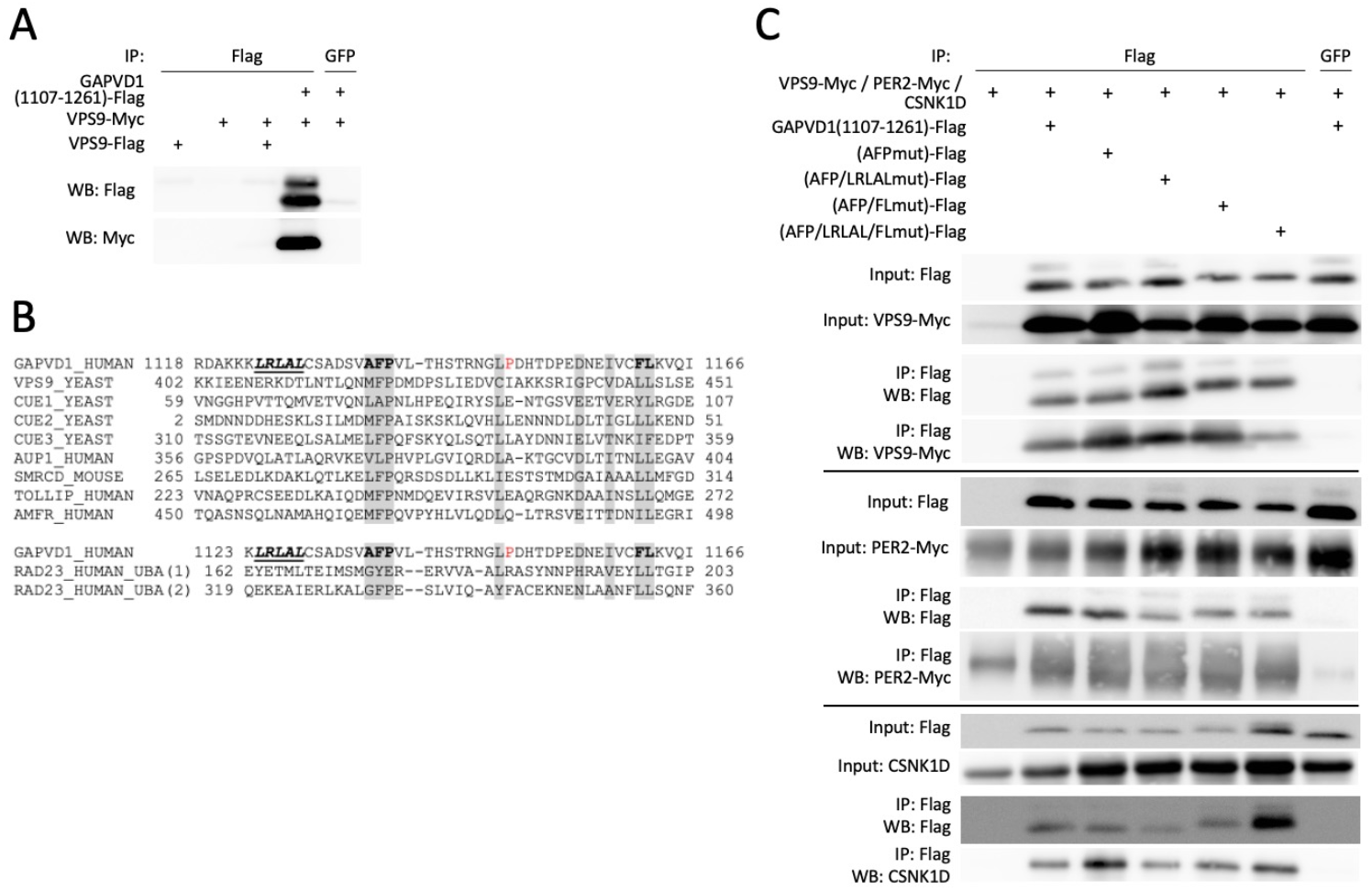
VPS9-interaction and CUE/UBA-like motifs in the GAPVD1 helical bundle. (**A**) VPS9 (GAPVD1(1261-1478)) and GAPVD1(1107-1261) were expressed in various combinations in BY-2 lysates and subjected to anti-Flag or control (anti-GFP) immunoprecipitation, followed by Western blotting with anti-Flag and anti-Myc antibodies. (**B**) The GAPVD1 helical bundle was aligned with representative CUE and UBA domains, highlighting conserved hydrophobic and acidic motifs at positions compatible with canonical ubiquitin-binding interfaces. Putative CUE/UBA core motifs are grey-boxed, a unique proline is marked in red, and an inverse ubiquitin-interacting motif (IUIM) is shown in underlined italics. Residues mutated in subsequent constructs are indicated in bold. (**C**) Flag-tagged helical bundle variants carrying combined mutations of putative CUE/UBA core residues and an N-terminal inverse ubiquitin-interacting motif, or lacking all motifs, were co-expressed with VPS9, PER2 or CSNK1D in BY-2 lysates and analyzed by anti-Flag immunoprecipitation and Western blotting. High background signal was observed for PER2-Myc in combination with Flag-beads.

### The GAPVD1 helical bundle is a non-canonical CUE/UBA-like protein-protein interaction domain

Sequence analysis of the helical bundle also revealed local homology to small helical ubiquitin-binding modules of the CUE (Coupling of ubiquitin conjugation to ER degradation) and UBA (Ubiquitin-associated) families, which typically comprise compact three-helix folds that recognize the Ile44-centered hydrophobic patch on ubiquitin and recruit their host proteins to ubiquitylated partners or membranes (**Fig. 2B**). Pertinent to the domain architecture of GAPVD1, CUE domains are frequently found adjacent to VPS9 domains in endosomal Rab-GEFs where they help couple Rab activation to ubiquitin-dependent cargo sorting (*15*). In the GAPVD1 helical bundle, conserved hydrophobic and acidic motifs occur at positions consistent with such interfaces, opening the possibility that the GAPVD1 helical bundle might function as a ubiquitin-directed localization or interaction module in certain cellular contexts. However, combined mutation of predicted CUE/UBA core motifs or an immediately N-terminal inverse ubiquitin-interacting motif (IUIM) did not impair binding of GAPVD1 to VPS9, PER2 or CSNK1D, and even a variant lacking all motifs bound as efficiently as the wild-type helical bundle (**Fig. 2C**). In line with these results, helix-propensity calculations place the second helix of the putative CUE/UBA-like region below the threshold for stable helix formation (*16, 13*). This marginal stability might further be affected by the presence of a proline residue toward the C-terminus of this helix, which is absent from the aligned canonical CUE/UBA sequences. Nevertheless, phosphorylation could, in principle, increase local helical propensity, and interaction with binding partners has been shown to promote helix formation in intrinsically disordered segments (*17*). Thus, despite its sequence similarity and predicted architecture, the GAPVD1 helical bundle does not behave as a canonical CUE domain but instead acts as a protein-protein interaction module that links GAPVD1 to PER2 and CSNK1D and is subject to intramolecular regulation of PER2-binding by the VPS9 domain.

### The PER2 casein kinase binding domain interacts with GAPVD1

Next, a set of C-terminal PER2 truncations was created, and binding of each variant to GAPVD1(680-1261) was analysed by co-immunoprecipitation. Variants with stepwise deletions of the CRY-binding domain (1-1148), the IDR (1-808), the casein kinase binding domain (CKBD) (1-476), and the PAS-B domain (1-323) revealed that binding of PER2 to GAPVD1 was lost in PER2(1-476) and therefore requires the CKBD (**Fig. 3A**). The identity of the CKBD was independently confirmed by co-immunoprecipitation of CSNK1D with the same set of PER2 truncation mutants (**Fig. 3B**). The requirement for the CKBD in PER2 for GAPVD1-binding raised the question if GAPVD1 and CSNK1D use the exact same protein surface in PER2 for interaction. However, the PER2 point mutants VL729-730GG and L730G, which had been shown previously to selectively interfere with CSNK1D-binding (*18*), can bind to full-length GAPVD1 and GAPVD1(680-1261), while immunoprecipitation of PER2-GFP and the PER2 point mutants confirmed that CSNK1D-binding is impaired in the PER2 variants VL729-730GG and L730G (**Fig. 3C**). Analogous results were obtained when PER2 and CSNK1D were expressed in BY-2 cell lysates. Full-length GAPVD1 and the minimal PER2-binding fragment GAPVD1(1107-1261) co-immunoprecipitated with both PER2(VL729-730GG) and PER2(L730G) (**Fig. 3D**). As a control, the lack of binding of PER2(VL729-730GG) to CSNK1D was confirmed in the cell-free system (**Fig. 3E**). Taken together, the GAPVD1-PER2 interaction is mediated by a helical bundle domain within amino acids 1114-1209 in GAPVD1 and the CKBD in PER2, whereby GAPVD1 binds to an interaction surface in the CKBD that is different from the one involved in CSNK1D binding.

**Figure 3.**
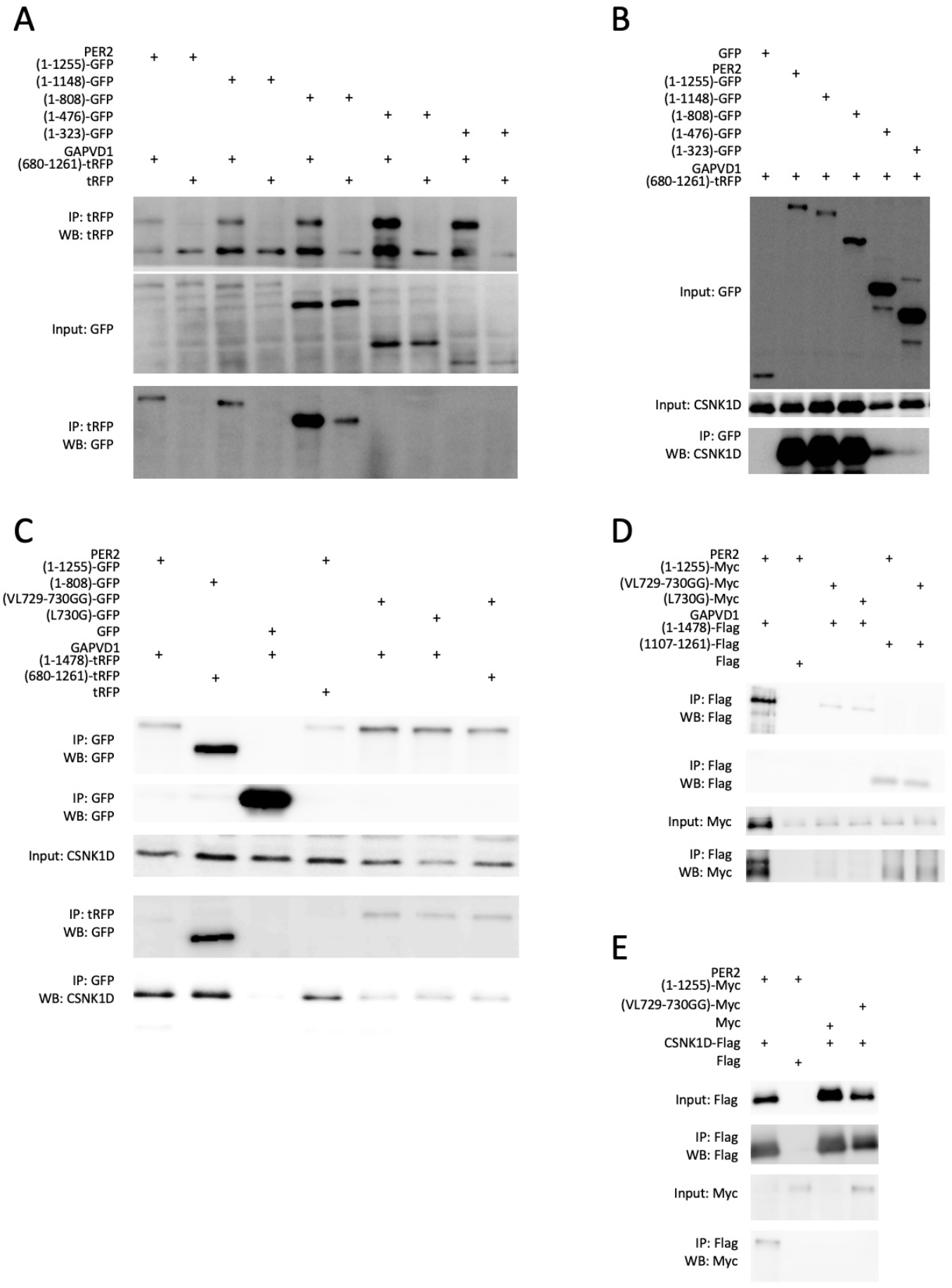
Mapping of the casein kinase binding domain in PER2 as the GAPVD1 interaction surface. (**A**) GFP-tagged PER2 full-length and C-terminal truncation constructs were expressed in HEK293T cells together with GAPVD1(680-1261)-tRFP. GFP IPs and input lysates were analyzed by immunoblotting for GFP and tRFP. (**B**) The same PER2-GFP truncation variants were expressed in HEK293T cells, and GFP IPs and inputs were probed for GFP and CSNK1D. (**C**) HEK293T cells were transfected with PER2-GFP constructs (1-1255 and mutants VL729-730GG, L730G) together with full-length GAPVD1(1-1478)-tRFP or GAPVD1(680-1261)-tRFP; GFP IP, tRFP IP and immunoblotting were used to detect PER2, GAPVD1, and CSNK1D. (**D**) In BY-2 lysates, PER2(1-1255)-Myc or PER2 mutants (VL729-730GG, L730G) were co-expressed with GAPVD1(1-1478)-Flag or GAPVD1(1107-1261)-Flag; anti-Flag IPs and inputs were analyzed by immunoblotting for Flag and Myc. (**E**) BY-2 lysates expressing CSNK1D-Flag with PER2(1-1255) Myc or PER2(VL729-730GG)-Myc were subjected to anti-Flag IP and immunoblotting for Flag and Myc. In some experiments, input samples could not be shown due to low protein abundance or antibodies unsuitable for input detection; in these cases, a direct immunoprecipitation is displayed instead.

### GAPVD1 attenuates PER2-CSNK1D binding and PER2 FASP phosphorylation

Interaction of GAPVD1 within the CKBD of PER2 hinted at a role for GAPVD1 in regulating PER2-CSNK1D binding. We addressed binding of PER2 to CSNK1D in GAPVD1 overexpressing HT1080 cells and HeLa *Gapvd1*^*-/-*^ knockout cells. Normalized to the amount of immunoprecipitated CSNK1D, endogenous PER2 was consistently less co-immunoprecipitated with CSNK1D when GAPVD1 was stably overexpressed in HT1080 cells (**Fig. 4A-B**). In line with this finding, more PER2 was bound to CSNK1D in *Gapvd1*^*-/-*^ HeLa cells compared to wild type cells (**Fig. 4A and C**). It was concluded that GAPVD1 negatively regulates the interaction between PER2 and CSNK1D, potentially modulating CSNK1D-mediated phosphorylation activity toward PER2. Indeed, full-length GAPVD1 and not GAPVD1(1107-1261) was able to suppress FASP phosphorylation of PER2 in HEK293T cells (**Fig. 4D**) and in an in vitro phosphorylation reaction with proteins expressed in BY-2 extracts (**Fig. 4E**). The selective inhibition by full-length GAPVD1 suggests that regions outside the minimal PER2/CSNK1D-binding fragment of GAPVD1 might be required to induce a CSNK1D-PER2 regulatory state that disfavours FASP modification.

**Figure 4.**
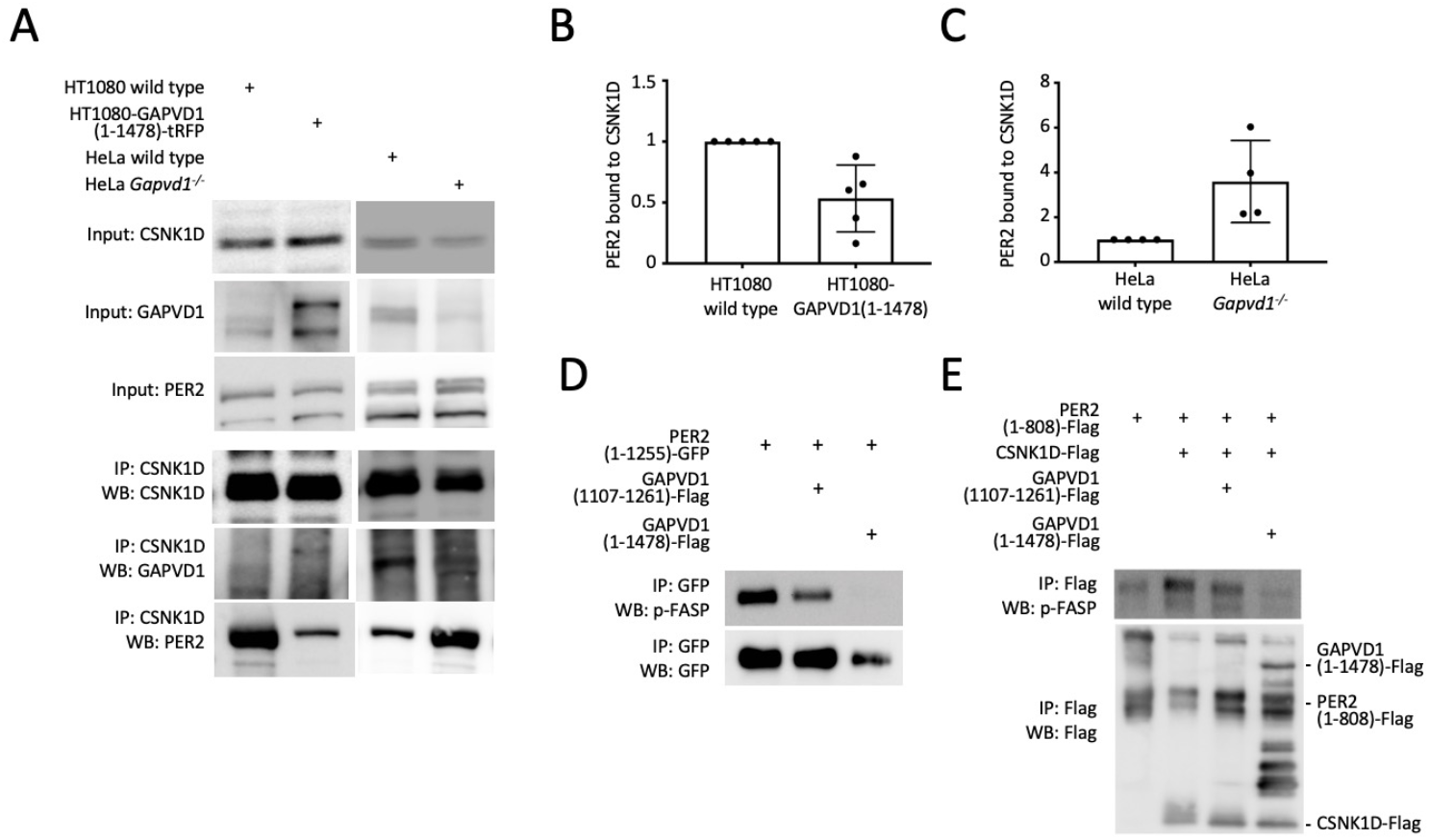
GAPVD1 attenuates PER2-CSNK1D association and PER2 FASP phosphorylation. (**A**) Endogenous CSNK1D was immunoprecipitated from HT1080 wild-type cells and cells stably expressing GAPVD1(1-1478)-tRFP, as well as HeLa wild-type and *Gapvd1*^*-/-*^ cells. IPs and inputs were analyzed by immunoblotting for CSNK1D, PER2 and GAPVD1-tRFP. (**B**) Band intensities of PER2 and CSNK1D from the IP samples in HT1080 cells were quantified by densitometry, and PER2 signal was normalized to immunoprecipitated CSNK1D for graphical display. (**C**) Densitometric analysis and normalization of PER2 relative to CSNK1D IP in HeLa wild-type and *Gapvd1*^*-/-*^ cells were performed and plotted in the same manner. (**D**) HEK293T cells were transiently transfected with PER2(1-1255)-GFP together with either GAPVD1(1107-1261)-Flag, or full-length GAPVD1(1-1478)-Flag. PER2(1-1255)-GFP was immunoprecipitated using anti-GFP antibodies, and IPs were analyzed by Western blotting with anti-GFP for total PER2 and a phospho-specific antibody directed against the PER2 FASP site. (**E**) In the Tobacco BY-2 cell-free expression system, PER2(1-808), CSNK1D, GAPVD1(1-1478) and GAPVD1(1107-1261) were expressed separately as Flag-tagged proteins and subsequently mixed and subjected to an in vitro phosphorylation reaction. Proteins were immunoprecipitated using anti-Flag, and IPs and inputs were probed by Western blotting with antibodies against Flag and the PER2 FASP phospho-specific epitope. In some experiments, input samples could not be shown due to low protein abundance or antibodies unsuitable for input detection; in these cases, a direct immunoprecipitation is displayed instead.

### Phosphorylation regulation of GAPVD1-PER2-CSNK1D complexes

Based on the observation that GAPVD1 affects PER2-CSNK1D binding and PER2 phosphorylation, we next asked whether GAPVD1 selectively attenuates PER2 FASP phosphorylation or reshapes global phosphorylation at the Degron/FASP region within PER2-CSNK1D complexes. To this end, we expressed PER2, GAPVD1, and CSNK1D in BY-2 lysates, performed in vitro kinase reactions, and analysed phosphopeptides by mass spectrometry. In PER2, numerous phosphosites within the Degron/FASP region were identified in samples containing CSNK1D and PER2, but none showed significant changes upon addition of GAPVD1 variants to the phosphorylation reaction **(Fig. 5A)**. Unfortunately, peptides spanning the FASP site were not detected, most likely due to technical limitations of peptide coverage and ionization in this region, a challenge also noted in a previous PER2 mass-spectrometry study that failed to robustly identify FASP-region phosphopeptides despite high overall sequence coverage (*19*). These data are therefore consistent with a scenario in which GAPVD1-dependent changes at the FASP region remain below the detection threshold in this assay, even though they are prominent when FASP phosphorylation is analysed with a specific antibody **(Fig. 4E)**.

**Figure 5.**
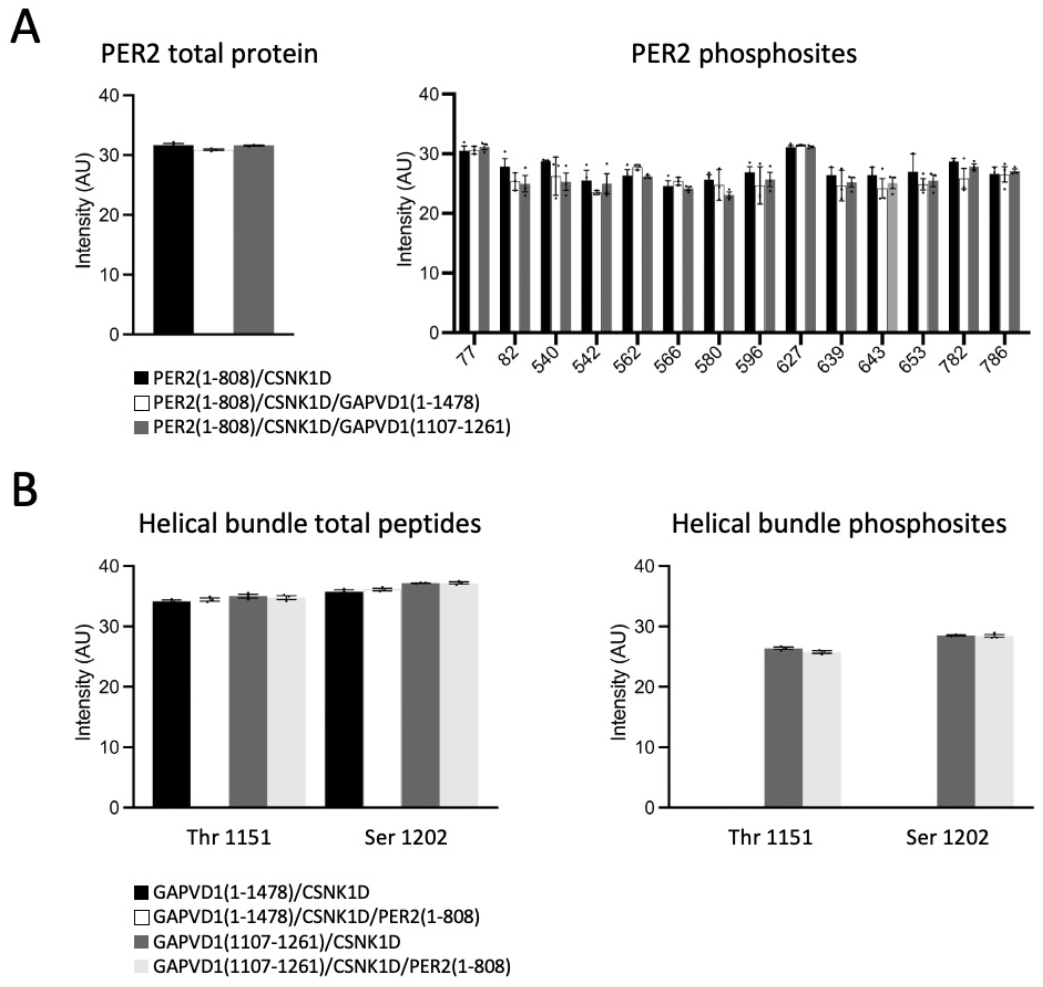
Phosphorylation analysis of PER-CSNK1D-GAPVD1 complex proteins. CSNK1D-Flag, PER2(1-808)-Flag, GAPVD1(1-1478)-Flag, or GAPVD1(1107-1261)-Flag were expressed separately in Tobacco BY-2 lysates, mixed, and subjected to anti-Flag immunoprecipitation after in vitro phosphorylation. Flag IPs were analyzed by LC– MS/MS to identify and quantify total protein or peptides and phosphopeptides of PER2 (**A**) and GAPVD1 (**B**).

When we examined GAPVD1 in these samples, we noticed that the only two peptides within the helical bundle domain that were reliably detected were only phosphorylated when this domain was expressed in isolation, whereas the same sites were not phosphorylated in the context of full-length GAPVD1, implying that surrounding regions or the higher-order architecture of the full-length protein restrict phosphorylation of specific protein residues while leaving them accessible in the isolated fragment **(Fig. 5B)**. Together with our earlier finding that the helical bundle can bind and stabilize the VPS9 domain, the selective phosphorylation of this region in isolation strengthens the idea that flanking segments regulate access to the helical bundle surface. Although deletion of flanking regions only enhanced access of PER2 **(Fig. 1C-E)** we have shown in a previous study that PER2 enhances CSNK1D-dependent GAPVD1 phosphorylation (*11*), suggesting a model in which PER2 binding and the surrounding regions of GAPVD1 might jointly regulate CSNK1D-dependent phosphorylation of the helical bundle. Overall, we conclude that GAPVD1’s main effect on PER2 is to modulate CSNK1D-dependent phosphorylation of the FASP site rather than to broadly remodel PER2 phosphorylation.

### Tail-independent regulation of PER2 FASP phosphorylation by CSNK1D

One important regulatory feature of CSNK1D in the PER2 phosphoswitch is its intrinsically disordered C-terminal tail, whose autophosphorylation can modulate kinase activity, substrate preference, and stability (*20, 8*). To test whether CSNK1D tail phosphorylation contributes to the mechanism that GAPVD1 employs to regulate PER2 FASP phosphorylation, we first sought to define how GAPVD1 affects CSNK1D autophosphorylation and kinase stability, and then asked whether these tail-dependent changes translate into altered PER2 modification. Attempts to quantify CSNK1D phosphorylation in the same BY-2 samples by mass spectrometry were inconclusive because phosphosite intensities varied strongly between replicates, precluding a reliable assessment of GAPVD1- or PER2-dependent changes in kinase autophosphorylation. We therefore turned to cell-based and targeted biochemical assays to probe CSNK1D stability and tail phosphorylation. CSNK1D protein levels differed between wild-type and *Gapvd1*^*-/-*^ cells, and treatment with the CSNK1D/E-inhibitor PF-670462 increased CSNK1D abundance in both backgrounds to a similar level **(Fig. 6A)**, consistent with the idea that GAPVD1-dependent differences in CSNK1D stability could partly arise from altered kinase phosphorylation, most likely at its autophosphorylated C-terminal tail (also compare levels of CSNK1D in GAPVD1-overexpressing and *GAPVD1*^*-/-*^ cells in **Fig. 4A)**. The impact of GAPVD1 on CSNK1D tail phosphorylation was analysed in HT1080 cells that stably express CSNK1D with or without GAPVD1. Expression of GAPVD1 robustly inhibited tail autophosphorylation of overexpressed and endogenous CSNK1D **(Fig. 6B)**. Similar results were observed upon expression of GAPVD1(1-1478) or GAPVD1(1107-1261) together with CSNK1D in vitro **(Fig. 6C)**. Interestingly, while GAPVD1(1107-1261) only weakly attenuated FASP phosphorylation compared to full-length GAPVD1 **(Fig. 4E)**, it efficiently suppressed CSNK1D tail phosphorylation **(Fig. 6C)**, indicating that GAPVD1 affects tail autophosphorylation and FASP site phosphorylation by independent mechanisms. Indeed, reduction of PER2 phosphorylation by GAPVD1 expression was also observed in a CSNK1D mutant lacking the C-terminal tail **(Fig. 6D)**. GAPVD1-mediated regulation of PER2 FASP phosphorylation is therefore not a consequence of altered CSNK1D tail phosphorylation because it does not require the tail at all. Collectively, these findings show that GAPVD1 modulates PER2 phosphorylation through regulation of PER2-CSNK1D binding and tail-independent inhibition of CSNK1D kinase activity.

**Figure 6.**
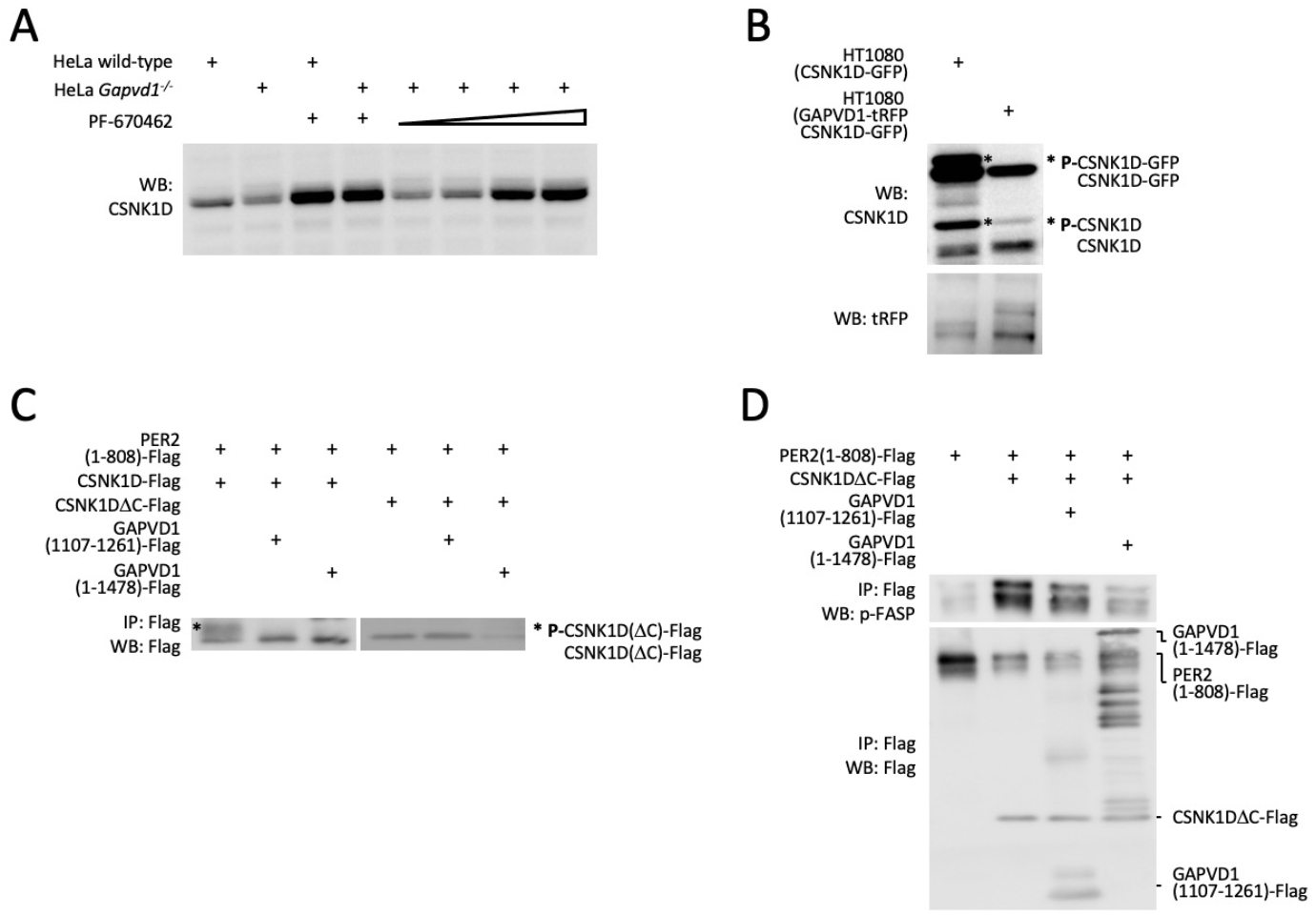
GAPVD1 regulates PER2 FASP phosphorylation independently from CSNK1D tail phosphorylation. (**A**) HeLa wild-type and *Gapvd1*^*-/-*^ cells were treated with PF-670462 or vehicle and analyzed by Western blotting using antibodies against CSNK1D to assess total kinase levels under basal and inhibitor-treated conditions. (**B**) HT1080 cells stably expressing CSNK1D-GFP with or without GAPVD1(1-1478)-tRFP were analyzed by Western blotting with anti-CSNK1D and anti-tRFP for GAPVD1. (**C**) In the Tobacco BY-2 cell-free expression system, CSNK1D-Flag, the C-terminal tail deletion mutant CSNK1DΔC-Flag, full-length GAPVD1(1-1478)-Flag, and GAPVD1(1107-1261)-Flag were expressed separately as Flag-tagged proteins and subsequently mixed and subjected to an in vitro phosphorylation reaction. Proteins were immunoprecipitated using anti-Flag, and IPs and inputs were probed by Western blotting with antibodies against Flag. (**D**) Similarly, CSNK1DΔC-Flag, PER2(1-808)-Flag, and GAPVD1 constructs (full-length or 1107-1261) were expressed separately as Flag-tagged proteins and subsequently mixed and subjected to an in vitro phosphorylation reaction. Proteins were immunoprecipitated using anti-Flag, and IPs and inputs were probed by Western blotting with antibodies against Flag and the PER2 FASP phospho-specific antibody.

## Discussion

GAPVD1 emerges from this study as a PER-complex scaffold that binds the PER2 CKBD via a compact helical bundle, engages CSNK1D independently with the same binding domain, weakens the PER2-CSNK1D interaction, and selectively inhibits CSNK1D activity toward the PER2 FASP site in a manner independent of the kinase’s C-terminal tail. GAPVD1 thereby acts as a novel regulator of the PER2 phosphoswitch within the PER2-CSNK1D control node. The helical bundle preceding the GAPVD1 VPS9 domain is sufficient for binding to PER2 and CSNK1D, whereas other parts of the protein including the catalytic RasGAP and VPS9 domains do not contribute detectably to these interactions and in fact attenuate the apparent affinity of the isolated helical bundle for PER2 when present in cis. GAPVD1 engages the PER2 CKBD on a surface that remains accessible in CKBD point mutants that abolish CSNK1D docking, demonstrating that GAPVD1 and CSNK1D contact overlapping yet distinct regions of the CKBD. Consistent with a direct binding mechanism, GAPVD1, PER2, and CSNK1D assemble into complexes and elicit distinct changes in their phosphorylation states when expressed in a heterologous cell-free extract that lacks other mammalian PER-complex components, supporting the view that their interactions are largely direct rather than scaffolded by additional protein factors. Deletion analyses further show that removing the VPS9 domain from otherwise binding-competent GAPVD1 variants markedly strengthens PER2 association without equivalently enhancing CSNK1D binding, indicating that VPS9 exerts a negative, PER2-selective regulatory influence on the helical bundle interaction surface. Consistent with this, the isolated helical bundle fragment is robustly phosphorylated by CSNK1D in vitro whereas the same region remains largely unphosphorylated in full-length GAPVD1, and the helical bundle can directly bind and stabilize VPS9. Together, these findings support a model in which intramolecular VPS9-helical bundle contacts shield parts of the docking and phosphorylation interface. Although sequence analyses revealed CUE/UBA-like motifs within the helical bundle, mutating these elements did not impair binding to PER2, CSNK1D, or VPS9, arguing that the helical bundle acts as a dedicated protein-protein interaction module rather than a canonical CUE/UBA ubiquitin-binding domain. Taken together, the finding that GAPVD1 attenuates PER2-CSNK1D binding and FASP-phosphorylation indicates that GAPVD1 modulates the PER2-CSNK1D interaction by effectively shortening CSNK1D dwell time on PER2 without fully disrupting CKBD anchoring, while internal interactions of the helical bundle with flanking GAPVD1 regions may provide an additional layer of regulation by dynamically modifying accessibility of the binding interface to PER2.

A central feature of the PER2 phosphoswitch is that stabilizing FASP phosphorylation antagonizes Degron phosphorylation and BTRC-dependent degradation, whereas enhanced Degron phosphorylation destabilizes PER2 and shortens circadian period length (*5*). The findings of this study show that full-length GAPVD1, but not a minimal CKBD-binding fragment, robustly reduces PER2 FASP phosphorylation, while mass spectrometry detects no global reductions in overall PER2 phosphorylation, with FASP and Degron sites not directly observed. Taken together with in vivo data, which indicate that Degron mutations can lengthen period and that Degron phosphorylation is the dominant driver of PER2 degradation (*21*), a likely model is that GAPVD1 primarily weakens the FASP “degradation brake” function rather than simply shutting off all CSNK1D activity toward PER2. This view is further supported by the observation that GAPVD1 knockdown lengthens not shortens the free-running period in fibroblasts (*9*), consistent with loss of a factor that normally pushes PER2 away from the stabilizing FASP arm of the phosphoswitch. Work on CSNK1D splice isoforms shows that tissue-specific expression of CSNK1D1 versus the more FASP-priming CSNK1D2, and activation-loop conformations that favour either the FASP or Degron site, can shift the balance of PER2 phosphorylation between stabilizing and degradative arms in a cell-type-dependent manner (*20, 22*). In such a model, GAPVD1-mediated weakening of FASP phosphorylation would push the phosphoswitch toward enhanced Degron activity or alternative degradation pathways, potentially shortening PER2 half-life in contexts where GAPVD1 is abundant and CSNK1D activity is already poised toward Degron use, whereas in cells where CSNK1D2-driven FASP priming predominates, GAPVD1’s impact might remain below the threshold required to measurably alter behavioural period. In agreement with this idea we observed that GAPVD1 knockout fails to alter circadian period, phase shifting properties or temperature compensation in U2OS cells (data not shown), despite the lengthened circadian period in GAPVD1-deficient mouse fibroblasts (*9*) and clear biochemical evidence from this study that GAPVD1 can modulate PER2 phosphorylation. A similar picture emerged from mechanistic work on the PER2 phosphoswitch in HEK293 cells, where CSNK1’s intrinsic preference for the stabilizing FASP region effectively limits usage of the canonical D2 Degron. At baseline, mutating D2 alone has essentially no effect on PER2 half-life, whereas preventing FASP phosphorylation shortens the half-life drastically (*21*).

Another key result is the persistent reduction of PER2 FASP phosphorylation by GAPVD1 when acting on a CSNK1D mutant lacking the C-terminal tail. Moreover, the minimal GAPVD1 binding fragment, although sufficient to block tail autophosphorylation, only weakly attenuates FASP phosphorylation compared to full-length GAPVD1. This decoupling implies that GAPVD1 regulates CSNK1D on at least two mechanistic layers: One, through reducing tail autophosphorylation, which might stabilize CSNK1D in a particular substrate-specific state; and two, which is more pertinent to phosphoswitch regulation, through a tail-independent effect, that might allosterically alter activation-loop conformation or peptide engagement at the FASP cluster through substrate competition. Molecular dynamics and Markov state modelling have shown that CSNK1D activation-loop conformations dictate FASP vs Degron preference, and that various point mutations such as tau can invert this preference independently of the kinase’s tail (*23*). Our data are consistent with GAPVD1 shifting the activation-loop toward a conformation that disfavours efficient FASP priming in the presence or absence of autoinhibitory tail contacts. In synopsis, the finding that CSNK1D tail regulation and FASP-site inhibition are mechanistically separable outputs of GAPVD1 binding is compatible with the idea that the helical bundle alone might stabilize a tail-engaging, autoinhibited conformation of CSNK1D without strongly perturbing PER2 docking or active-site access. In contrast, additional regions present in full-length GAPVD1 could be required either to remodel the PER2-CSNK1D interface or to present phosphorylated GAPVD1 segments as competing substrates that selectively disfavour FASP phosphorylation.

Overall, our previous and current data are consistent with a model centred around stoichiometry-dependent control of CSNK1D within the PER2-GAPVD1 module. In cells or subcellular compartments with high PER2 relative to GAPVD1, PER2 appears to preferentially access and activate tail-inhibited CSNK1D, thereby stimulating CSNK1D-dependent GAPVD1 phosphorylation and degradation (*11*). Conversely, when GAPVD1 is abundant, full-length GAPVD1 efficiently binds CSNK1D and the PER2 CKBD, weakens PER2-CSNK1D association and attenuates CSNK1D activity toward the PER2 FASP site. We speculate that multiple phosphorylated sites within the disordered GAPVD1 IDR could act as low-affinity phosphopeptide mimetics that transiently engage CSNK1D substrate-recognition surfaces, thereby competing with FASP for productive active-site access and further biasing the phosphoswitch toward reduced FASP phosphorylation. In this scenario the relative amounts and timing of PER2 and GAPVD1 would help determine which partner gains productive access to CSNK1D and whether its activity is directed more toward GAPVD1 phosphorylation or PER2 FASP phosphorylation. Direct quantitative measurements of binding affinities will be required to test this explicitly.

Future work should attempt to integrate GAPVD1 into quantitative phosphoswitch models. Existing models have successfully recapitulated period, temperature compensation, and responsiveness to kinase perturbations using CSNK1 kinetics, FASP/Degron hierarchies, and simple binding equilibria (*24*). Our data indicate that an additional state variable - interaction of GAPVD1 with PER2 and CSNK1D with concomitant phosphorylation inhibition - might be required to capture situations where FASP phosphorylation is selectively reduced without a proportional change in Degron phosphorylation or global CSNK1D activity. Such a model might be able to predict conditions under which changes in GAPVD1 expression or phosphorylation produce measurable changes in PER2 half-life and behavioural period, and when they are buffered by the intrinsic phosphoswitch design.

## Materials and Methods

### Cell culture and generation of stable lines

HEK293T, HeLa and HT1080 cells were maintained in Dulbecco’s modified Eagle’s medium (DMEM) supplemented with 10% fetal bovine serum, 1% penicillin/streptomycin and 2 mM L-glutamine at 37 °C in a humidified 5% CO_2_ incubator. U2OS PER2::LUC and BMAL1::LUC reporter cells were cultured under the same conditions. Stable HT1080 PER2-GFP, CSNK1D-GFP and GAPVD1-tRFP cell lines were generated by transfection with the respective expression plasmids followed by antibiotic selection and single-cell cloning as described previously for PER2-GFP-expressing HT1080 cells (*11*).

### CRISPR/Cas9-mediated knockout

U2OS reporter cell lines stably expressing PER2::LUC or BMAL1::LUC were used to generate GAPVD1 knockout derivatives. Cells were transfected with CRISPR/Cas9 constructs expressing Cas9 and a puromycin resistance cassette together with sgRNAs targeting human GAPVD1, using two independent single-guide RNAs per reporter line directed against different exons of the GAPVD1 coding sequence (GAPVD1 exon 2: forward 5′-GAGAATTCTTGAGTCGATTG-3′ and reverse 5′-CAATCGACTCAAGAATTCTC-3′; GAPVD1 exon 5: forward 5′-TTTATTGGCAAACCTACCCC-3′ and reverse 5′-GGGGTAGGTTTGCCAATAAA-3′). Mock-transfected cells, treated with transfection reagent and empty vector, were processed in parallel. Following puromycin selection and single-cell cloning, individual clones were expanded and screened for GAPVD1 loss by standard molecular and protein-based assays. For circadian phenotyping, parental and GAPVD1-deficient PER2::LUC and BMAL1::LUC cells, together with mock controls, were analyzed for free-running period length, responses to phase resetting by dexamethasone, and temperature compensation in bioluminescence recording assays under constant conditions.

### Plasmids and molecular cloning

Full-length human GAPVD1 was amplified from pCDNA3-2xHA-hRME-6 (*25*) using primers introducing MluI and ApaI sites and subcloned via pCR2.1-TOPO into the pMCC-tRFP-H backbone to generate GAPVD1-tRFP. GAPVD1 truncation and deletion constructs were generated by restriction digestion and ligation of short hybridized oligonucleotides or by overlap PCR followed by subcloning and transfer into pMCC-tRFP-H and pALiCE01. All PCR-derived regions and junctions were verified by Sanger sequencing.

### Transient transfection and pharmacological treatments

For transient expression experiments, mammalian cells were seeded one day before transfection to reach 60–80% confluence and transfected with the indicated plasmids using Effectene transfection reagent according to the manufacturer’s instructions. Plasmid amounts were adjusted so that the total DNA per dish remained constant by addition of empty vector. The cells were harvested after 48 hours and frozen at -80°C for later use. Where indicated, cells were treated with the CSNK1D/E inhibitor PF-670462, which was added from a DMSO stock solution to the culture medium at the specified final concentration; control cells received an equivalent volume of DMSO.

### Co-immunoprecipitation and immunoblotting

#### Immunoprecipitation of proteins using GFP nanobodies coupled to agarose beads

Cells were washed with ice-cold PBS and lysed in CHAPS lysis buffer (20 mM Tris HCl pH 7.5, 150 mM NaCl, 0.03% CHAPS, 0.5 mM EDTA) supplemented with PhosStop EASYpack and protease inhibitor cocktail (Roche, Switzerland). Insoluble material was removed by centrifugation, and cleared lysates were incubated with GFP/SPOT:SSSSS-Trap® beads (ChromoTek) overnight at 4 °C. After washing in high salt CHAPS lysis buffer (20 mM Tris HCl pH 7.5, 500 mM NaCl, 0.03% CHAPS, 0.5 mM EDTA) and low salt CHAPS lysis buffer (20 mM Tris HCl pH 7.5, 0.03% CHAPS, 0.5 mM EDTA) bound proteins were eluted in Laemmli sample buffer, separated by SDS-PAGE and transferred to PVDF membranes.

#### Immunoprecipitation of proteins using magnetic microbeads

Cells were washed with ice-cold PBS and lysed in Triton X-100 lysis buffer (150 mM NaCl, 1% Triton X-100, 50 mM Tris HCl pH 8.0, PhosStop and protease inhibitor). Insoluble material was removed by centrifugation and cleared lysates were incubated with protein A or G (Miltenyi Biotec) and 4 µg tRFP, GAPVD1 or CSNK1D antibodies at 4°C overnight. Cell lysates were loaded onto equilibrated μColumns (Miltenyi Biotec) in the μMACS™ Separator magnetic field. Columns were washed with high salt lysis buffer (500 mM NaCl, 1% NP-40, 50 mM Tris HCl pH 8.0) and low salt lysis buffer (1% NP-40, 50 mM Tris HCl pH 8.0). Proteins were eluted in Laemmli sample buffer, separated by SDS-PAGE and transferred to PVDF membranes.

#### Antibodies for immunoprecipitation and western blot analysis

GAPVD1 (#NBP1-19156, Novus Biologicals), PER2 (#20359-1-AP, Proteintech), CSNK1D (#AB85320, Abcam), GFP (#632381, Clontech), TagRFP/tRFP (R10367 and MA5-15257, Thermo Fisher Scientific / #AB233, Evrogen), Flag (F1804, Sigma-Aldrich), Myc (M5546, Merck/Sigma-Aldrich), Spot (28a5, ChromoTek/Proteintech), and phospho-FASP PER2 (#EPR19820, Abcam).

### Tobacco BY-2 cell-free expression and in vitro phosphorylation

Plasmid DNA encoding GAPVD1, PER2 and CSNK1D variants were used at 5 nM with 50 μL ALiCE® reaction mix (LenioBio). Reactions were incubated in an orbital shaker at 700 rpm and 25°C for 48 h. Synthesized proteins were subject to immunoprecipitation or used for in vitro kinase reactions for 6 h in 50 mM Tris-Cl pH 7.3, 50 mM KCl, 10 mM MgCl_2_, 20 mM β-glycerophosphate, 15 mM EGTA, 2 mM ATP followed by immunoprecipitation, western blot, or mass spectrometry analysis.

### Mass spectrometry-based phosphosite analysis

Samples from the tobacco cell lysate were separated by SDS-PAGE and three slices per gel lane were excised. Tryptic protein digestion and peptide purification was carried out as described previously (*26*). Prior to LC-MS analysis the peptides were resuspended in 3% formic acid (FA)/1% acetonitrile (ACN) and loaded onto a nanoLC system (RSLCnano, Thermo Scientific). The peptides were first trapped on a trapping column (Acclaim PepMap100, C18, 5 µm, 100 Å, 300 µm i.d. × 5 mm, Thermo Scientific) for 10 minutes. Subsequently, the peptides were separated on an analytical column (Aurora Ultimate 25*75 C18 UHPL column, Ionopticks; 45°C column oven temperature) for 40 minutes. Gradient settings: 0–2 min: 1-2 % buffer B (buffer A: 0.1 % FA; buffer B: 80 % ACN, 0.1 % FA), 2-29 min: 2–50 % buffer B, 29-31 min: 50-99 % buffer B, 31-33 min: 99 % buffer B, 33-35 min: 99-2 % buffer B, 35-40 min: 2 % buffer B.

Data acquisition was performed on an Exploris 480 mass spectrometer (Thermo Scientific) in data-dependent acquisition (DDA) mode. MS Settings: Resolution 60k; scan range: 350-1250; AGC target: standard; max injection time mode: Auto; MS2: Resolution 15k; normalized HCD collision energy: 30%; AGC target: 100%; max injection time: 55 ms. Charge state: 2-5.

The raw data was analyzed using MaxQuant (v 2.6.3.0) against a combined *Nicotiana tabacum/Homo sapiens* database (07/2023) with the individual sequences of the recombinant proteins added using the Andromeda search engine with default mass tolerance settings. Trypsin was set as the protease (two allowed missed cleavages). Fixed modification: Carbamidomethylation (Cys); variable modifications: Oxidation (Met), Phosphorylation (Ser, Thr, Tyr) and N-terminal protein acetylation. False discovery rates: 0.01 for peptides, proteins and modification sites; minimum peptide score for modified peptides: 40; minimum peptide length: seven amino acids.

## Acknowledgments

We thank Kathy L. Gould (Vanderbilt University) for generously providing the GAPVD1 knockout cell lines used in this study and Christian Mielke (University Hospital Düsseldorf) for valuable help with the CRISPR/Cas methodology.

## Author contributions

Conceptualization: HI, FK, HR

Methodology: EW, CP

Investigation: HI, CB, YC, JD, LL, VO, SS, EW, HW, VZ

Visualization: HI, FK, HR

Supervision: FK, HR

Writing—original draft: HR

Writing—review & editing: HI, CP, FK, HR

## Data and materials availability

All data are available in the main text or the supplementary materials.

